# Reactivation of hedonic but not sensory representations in human emotional learning

**DOI:** 10.1101/2021.11.25.469891

**Authors:** M. R. Ehlers, J. H. Kryklywy, A. O. Beukers, S. R. Moore, B. J. Forys, A.K. Anderson, R. M. Todd

## Abstract

Learning which stimuli in our environment co-occur with painful or pleasurable events is critical for survival. Previous research has established the basic neural and behavioural mechanisms of aversive and appetitive conditioning; however, it is unclear what precisely is learned. Here we examined what aspects of the unconditioned stimulus (US) – sensory and hedonic – are transferred to the conditioned stimulus (CS). To decode the content of brain activation patterns elicited during appetitive (soft touch) and aversive (painful touch) conditioning of faces, a novel variation of representational similarity analysis (RSA) based on theoretically driven representational patterns of interest (POIs) was applied to fMRI data. Once face associations were learned through conditioning, globally the CS reactivated US representational patterns showing conditioning-dependent reactivation. More specifically, in higher order brain regions, the CS only reactivated hedonic but not sensory aspects of the US – suggesting that affective conditioning primarily carries forward the valence of the experience rather than its sensory origins.

## Introduction

The capacity to develop painful or pleasurable associations with predictive cues is highly conserved across species (Clark et al., 2002; Kryklywy et al., 2020; Pessoa et al., 2019). It is so central to our survival and our ability to make sense of the world that, often, a single exposure to an aversive or appetitive stimulus and an associated neutral stimulus is enough for us to remember the critical information for the rest of our lives (VanElzakker et al., 2014; Yamamoto et al., 1994). Decades of research on emotional learning processes have established basic neural and behavioral mechanisms by which human and non-human animals learn what cues (conditioned stimulus, CS+) predict the occurrence of inherently positive or negative events (unconditioned stimuli, US) (Andreatta & Pauli, 2015; Garcia & Koelling, 1966; LeDoux, 2003; Maren, 2001; Martin-Soelch et al., 2007). Yet longstanding fundamental questions about the nature of the information that we learn to associate with salient events remain to be resolved: When a conditioned cue is encountered, does the brain directly recapitulate representations of the pleasant or unpleasant experience the cue predicts? If so, what content is represented? Do objects associated with emotionally relevant events evoke the visceral sensory properties of the original salient event, or unconditioned stimulus? Or do they rather evoke the hedonic response to the experience, that is the painful and pleasurable qualities that were experienced along with the event?

These questions go back to different theories of learning. The stimulus-response (S-R) school argued that conditioning involves a representation of the association between a stimulus and its response (Holland, 2008), which can be a motor, physiological or hedonic response. In contrast, the stimulus-stimulus school (S-S) (Byrne, 2003) has proposed that, with learning, the initially neutral CS+ comes to elicit the same afferent activity initially elicited by the innately arousing US (Hull, 1943), and an association between a sensory process and a motor response is formed (Spence, 1950). These questions about the nature of emotional experience – about overlap and interaction between sensation and cognition – are rooted in antiquity. Classical philosophers have already debated whether emotion existed as a unitary experience, or as a set of dissociable constructs, with unique components of cognitive appraisal and biological drives (*Epicurus | Internet Encyclopedia of Philosophy*, n.d.; Sihvola & Engberg-Pedersen, 1998). Interpreted in the context of the current study, they argued whether being exposed to an appetitive or aversive event evokes a unitary experience involving both the activation of sensory organs and a hedonic appraisal of the situation, or whether sensation and cognition are two distinct entities independently contributing to an emotional experience that, in turn, can be independently transferred to a neutral stimulus.

Today, behavioral evidence supporting either argument can be found. For example, *autoshaping* or *sign-tracking* describes a phenomenon in classical conditioning in which animals behaviorally engage with the CS in the same way as they would with the US (Brown & Jenkins, 1968; Flagel et al., 2009). Critically, this effect is so far reaching that some animals completely neglect the US, thereby suggesting that S-S conditioning has occurred (Morrison et al., 2015). By contrast, *outcome devaluation* experiments related to habit formation suggest that once a certain action is established, a set response to a stimulus is initiated irrespective of the associated outcome, thereby indicating S-R learning has occurred (Schwabe et al., 2007; Smith & Graybiel, 2016). Yet, a lack of appropriate techniques has hampered investigations into neurobiological representations that could support our understanding of what is learned and transferred during affective conditioning.

Recent advancements in functional neuroimaging, however, are providing exciting new avenues towards addressing these questions. For example, in a series of experiments, Visser and colleagues (Visser et al., 2011, 2013, 2015) employed trial-by-trial representational similarity analysis (RSA), a multivariate approach to examining neural instantiations of the degree to which content is categorized as more or less similar, to examine how associative learning changes the representation of the initially neutral CS+. The authors showed that, before fear conditioning, neutral stimuli were represented as most similar to other stimuli with similar visual characteristics. However, after conditioning, stimuli paired with a US (the CS+), and those without pairing (the CS-) formed separate representational categories, irrespective of their visual similarity (Visser et al., 2011). The initial classification based on visual categories was overwritten by the emotional association. Another study compared the neural pattern activation in response to US and CS+ during a fear conditioning paradigm and tracked how the reactivation of the US pattern by the CS+ develops over the course of learning in the insula (Onat & Büchel, 2015). However, these studies did not probe the *content* of the reactivated patterns they observed; that is, they could not demonstrate if the US pattern gets reactivated in its entirety as opposed to only components of it. Thus, the goal of the present study was to build upon this work by investigating not only how the CS+ changes in representation following conditioning, but also whether a CS+ reactivates brain activation patterns elicited by the US both in its general representational form and as discriminable non-hedonic and hedonic components.

To do this, we employ Representational Similarity Analysis (RSA) (Kriegeskorte et al., 2008; Kriegeskorte & Kievit, 2013), a multivariate approach to functional neuroimaging analyses, that allows us to investigate population code representations. RSA was combined with a hypothesis-driven variation of Pattern Component Modeling (PCM) (Diedrichsen et al., 2018). This is an approach we recently developed to use fMRI to decode and fit representational patterns in response to appetitive and aversive US to predefined pattern components modeling theoretical representations patterns of interest (POIs) present in the US evoked signal (Kryklywy, Ehlers, et al., 2021). With this technique, we could investigate the extent to which hedonic and non-hedonic aspects of sensation are reproduced by the CS as novel affective associations are acquired.

Participants completed two conditioning paradigms in the MR scanner in which either an appetitive brush stroke to the forearm or painful pressure applied to the thumbnail were paired with the presentation of specific facial identities. Representational similarity patterns from eight regions of interest for CS and US were extracted on a trial-by-trial basis by means of RSA with theoretically based pre-specified patterns of interest (POIs) which we identified via PCM. We found that, once learned associations were established, the CS reactivates US representational patterns in brain regions typically associated with conditioning (e.g. amygdala, ventromedial prefrontal cortex and insula). By comparing patterns of representation with theoretically grounded POIs, we demonstrate that primary sensory regions reactivate components of the US that do not rely on learni ng. In contrast, higher order brain regions reactivate representations of hedonic value of the US, supporting a model of stimulus-response learning in the human brain.

## Results

In separate tasks for appetitive and aversive associative learning, two different faces (male and female for CS+ in aversive and appetitive tasks) were paired with either pleasurable brush strokes to the forearm or aversive pressure to the thumbnail (US). Two other faces (CS-) were never paired with the US (see Figure 1a). All faces wore affectively neutral expressions and individual identities were counterbalanced between participants.

**Figure 1.**
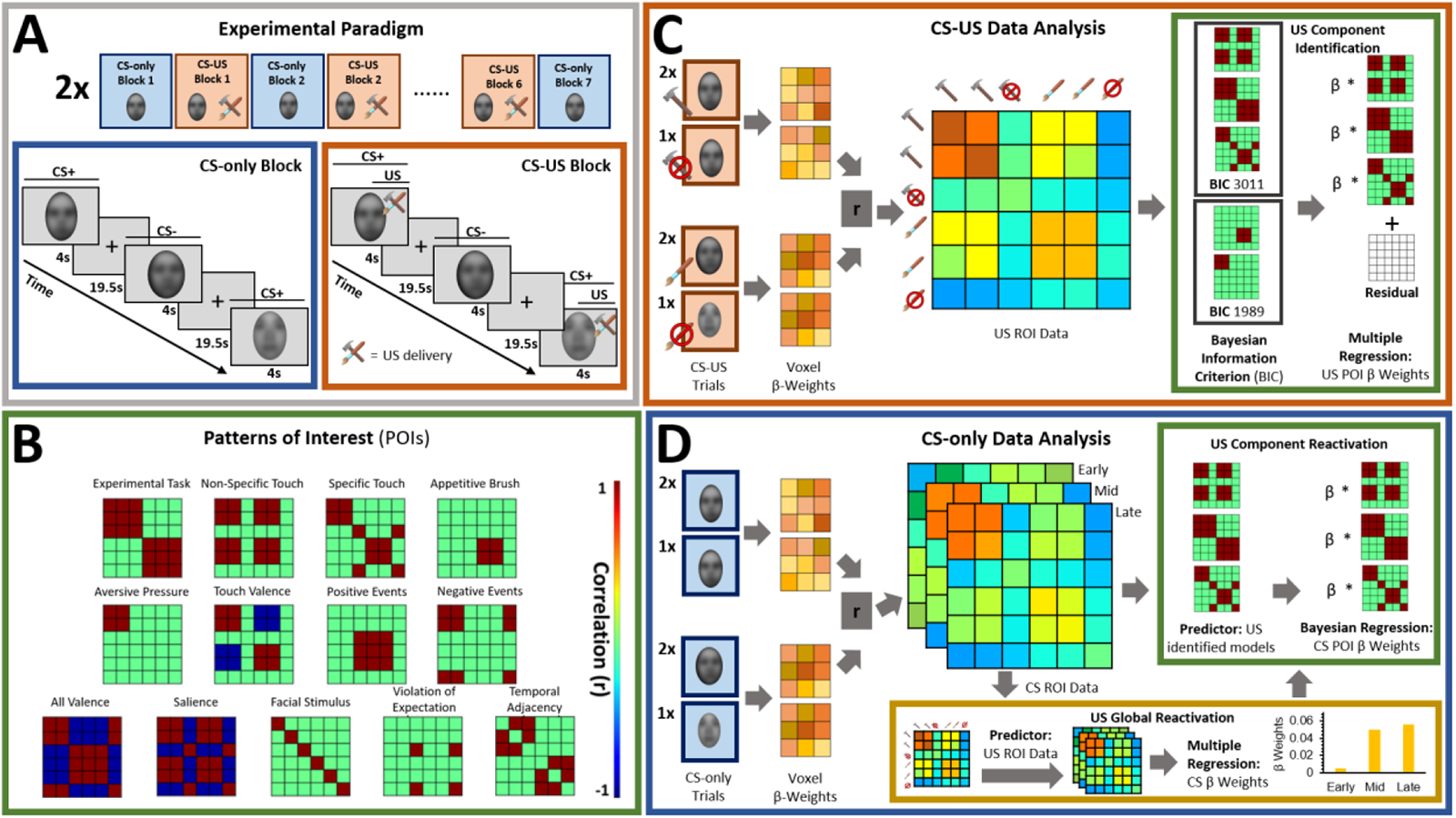
**A.** Both experimental tasks (appetitive and aversive conditioning) followed the same general structure in which 7 CS- only blocks were interleaved with 6 CS- US paired blocks. In CS- only blocks, faces (blurred on figure only) were presented by themselves, while the CS+ faces were paired with appetitive brush or aversive pressure in the CS- US blocks. **B.** Patterns of interest (POIs) demonstrate the representational pattern that would be observed in the experimental data if a region of interest (ROI) perfectly represented the theoretically derived constructs. **C.** Voxel activation patterns in each ROI, averaged over all CS- US trials for each task condition, were correlated with those in each other task condition using a representational similarity analysis (RSA) approach. Bayesian Information Criterion (BIC) was used to find the best combination of POIs for each ROI. Multiple regression was used to obtain beta coefficients for each POI (Kryklywy, Ehlers, et al., 2021). **D.** RSA was performed separately on early, mid, and late CS- only trials to examine reactivation of US information over the course of conditioning. (Bayesian) multiple regression was used in order to quantify the US model reactivation by CS- only data globally and BIC identified components were used to quantify which US aspects were reactivated by CS.

### Subjective ratings

For each face, ratings of likeability and trustworthiness were acquired before and after associative learning. CS discrimination, i.e. [CS+ minus CS-], in likeability and trustworthiness ratings was calculated for ratings obtained before and after completion of the appetitive and aversive tasks. Before appetitive conditioning, no CS discrimination was evident in likeability, *t*(68) = 0.29, *p* = .772, 95% CI [-3.66, 4.90], or trustworthiness ratings, *t*(68) = 1.21, *p* = .232, 95% CI [-1.21, 6.98] as expected for two neutral facial expressions. Likewise no difference between CS+ and CS- likeability, *t*(68) = -0.92, *p* = .359, 95% CI [-5.77, 2.12] and trustworthiness, *t*(68) = -1.52, *p* = .134, 95% CI [-6.93, 0.94] ratings was measured before aversive conditioning. Importantly however, CS+ and CS- likeability (appetitive conditioning: *t*(68) = -3.52, *p* < .001, 95% CI [-13.59, -3.76]; aversive conditioning: *t*(68) = 2.61, *p* = .011, 95% CI [1.89, 14.06]) differed significantly after conditioning such that in the appetitive task, the CS+ was more likeable, while in the aversive task, the CS- was more likeable indicating successful conditioning in both tasks. CS discrimination in trustworthiness reached significance only in the appetitive, *t*(68) = -2.58, *p* < .012, 95% CI [-11.47, -1.45], but not in the aversive task, *t*(68) = 1.45, *p* < .152, 95% CI [-0.94, 5.92]. Overall stimulus ratings indicate successful conditioning in both domains.

### US pattern reactivation by CS

The goal of the current study was to determine the extent and content of US pattern reactivation by the CS once associations were learned. This was achieved by comparing patterns of information representation identified during CS- only conditions early, mid, and late in conditioning to previously identified representational patterns observed in CS- US paired trials. The extent was assessed as both a global reactivation of US representational patterns, and the reactivation of specific informative patterns of interest (POIs, Figure 1b), which further provided information about the content of reactivation.

After initial preprocessing and first-level analysis of fMRI data, representational similarity analysis (RSA) was conducted on the patterns of BOLD activity within predefined regions of interest (ROIs). This resulted in a set of similarity matrices, comprised of correlations between all pairs of conditions (separate for CS- US paired, and CS- only trials; see Figure 1c/d). Representational information in BOLD responses to the US was identified by using Bayesian Information Criterion (BIC) to fit predefined POIs – idealized similarity matrices modelling specific information content (e.g., the valence of the touch stimulus) – to the observed US similarity data (Figure 1c). The representation of appetitive and aversive US, independent from CS- only trials, are focus of a different study (Kryklywy, Ehlers, et al., 2021).

To identify the extent to which US patterns were reactivated by CS following learning, a reconstructed measure of US activation similarity (rUS) was fit to CS similarity data from early-, mid-, and late-conditioning trials. rUS were constructed by summing the scaled contribution of each POI contributing to overall similarity in an ROI (identified though hypothesis-driven PCM; see (Kryklywy, Ehlers, et al., 2021). Next, to examine the degree to which each specific US-defined POI was comparable to the pattern of CS activation – the content driven reactivation – separate Bayesian linear models with the combination of US–defined POIs as predictors, and rUS and CS representational patterns as outcome variables were fitted in order to compare the (beta) weight of each predictor for the CS representational pattern to that of the US (see Figure 1d).

A reactivation of the following US-defined POIs is interpreted as reactivation of hedonic information: ‘Aversive Pressure’/‘Appetitive Brush’ (representation of either US type independently); ‘Touch Valence’ (representation of the positive relative to negative valence of US stimuli on a single continuum); and ‘Negative Events’ (representation of both aversive pressure and absence of pleasurable touch). Interpretation of these as hedonic information is particularly strong if these POIs are observed *without* reactivation of general ‘Non-Specific Touch’ representational patterns (representation of the tactile manipulation independent of affective discrimination).

#### Sensory regions of interest

##### Primary Somatosensory Cortex (S1)

Globally, the rUS pattern predicted the CS representation in mid and late conditioning, while the association was at trend level for early conditioning (*p* = .069) (see Table 1). Thus, there was some increase in the reliability of global reactivation effects with learning. The analysis of POI reactivation (See Figure 2) showed that the contribution of the POI ‘Experimental Task’, where there is similarity between CS+ and CS- within each conditioning task, to CS representational patterns was consistent with that of the US in mid and late conditioning, showing a conditioning effect. In contrast, ‘Non-specific Touch’ was consistent with the US pattern in early and late conditioning, a pattern inconsistent with the conditioning effect (see Figure 2).

**Table 1.** Global US reactivation by CS representation patterns in early, mid and late conditioning

Global reactivation (df = 1, 1489)
| ROI | Early Conditioning |  |  |  | Mid Conditioning |  |  |  | Late Conditioning |  |  |  |
| --- | --- | --- | --- | --- | --- | --- | --- | --- | --- | --- | --- | --- |
| | Adj.R<sup>2</sup> | F | p | $\beta$ | Adj.R<sup>2</sup> | F | p | $\beta$ | Adj.R<sup>2</sup> | F | p | $\beta$ |
| S1 | .002 | 3.311 | .069 | .185 | .004 | 6.671 | **.010** | .287 | .003 | 6.087 | **.014** | .281 |
| V1 | .003 | 6.082 | **.014** | .734 | .004 | 7.241 | **.007** | .842 | .007 | 10.55 | **.001** | 1.086 |
| VVS | .005 | 8.843 | **.003** | .812 | .005 | 8.766 | **.003** | .848 | .003 | 5.188 | **.023** | .713 |
| Amy | -.5.266e | 0.922 | .337 | -.175 | .0003 | 1.518 | .218 | .232 | .003 | 5.562 | **.018** | .471 |
| vmPFC | -.0003 | 0.487 | .486 | -.153 | .006 | 9.091 | **.003** | 0.743 | .005 | 8.537 | **.004** | .723 |
| ACC | .0006 | 1.937 | .164 | -.177 | .004 | 6.704 | **.010** | .357 | .011 | 17.20 | **<.001** | .609 |
| aIns | -.0001 | 0.792 | .374 | .101 | .011 | 17.18 | **<.001** | .515 | .015 | 23.13 | **<.001** | .630 |
| pIns | -.0006 | 0.018 | .893 | .010 | .006 | 10.43 | **.001** | .248 | .002 | 4.221 | **.040** | .173 |

S1 = primary somatosensory cortex; V1 = primary/secondary visual cortex; VVS = ventral visual structures; Amy = amygdala; vmPFC = ventromedial prefrontal cortex; ACC = anterior cingulate cortex; aIns = anterior insula; pIns = posterior insula; bolded p-value indicates significance at $\alpha = 0.05$

**Figure 2.**
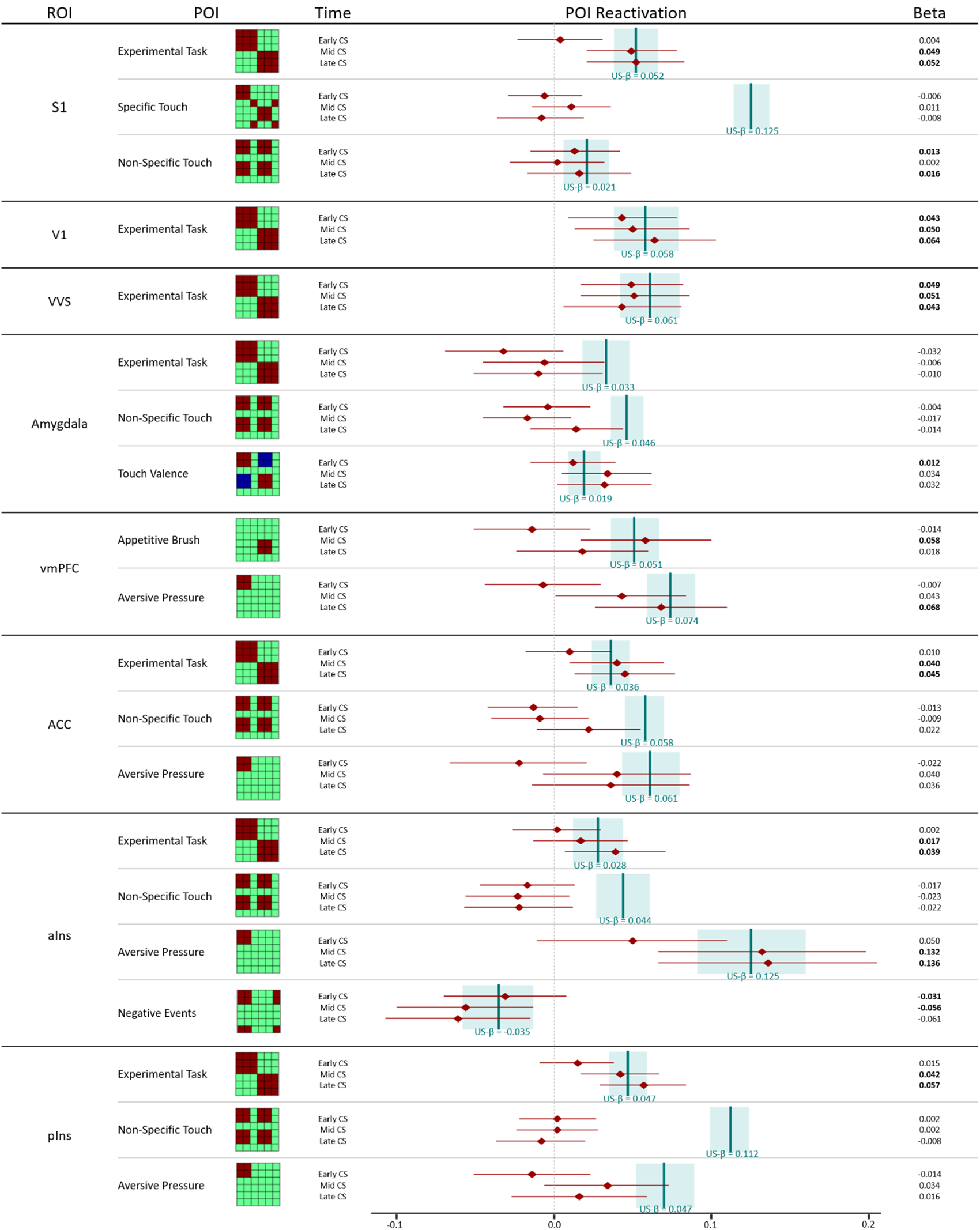
Pattern of Interest (POI) reactivation by reconstructed US (rUS) as well as early, mid and late CS data in different regions of interest (ROIs). Displayed are beta weights and 95% credible intervals for each POI reactivation obtained from separate Bayesian linear models performed on rUS and CS data with the illustrated POIs as predictors. The contribution of any given POI at any given time point of CS conditioning is considered consistent with that POIs contribution to rUS data if the point estimate of the CS data (in red) falls within the credible interval of the rUS data (in blue). Consistent contribution is further indicated by printing the obtained beta weight in bold. S1 = primary somatosensory cortex; V1 = primary/secondary visual cortex; VVS = ventral visual structures; vmPFC = ventromedial prefrontal cortex; ACC = anterior cingulate cortex; aIns = anterior insula; pIns = posterior insula

##### Primary/secondary visual cortex (V1)

Global CS representational patterns were significantly predicted by V1 rUS during early, mid, and late conditioning, indicating no effect of conditioning (see Table 1). Likewise, the reactivation of the POI ‘Experimental Task’ was consistent with a contribution to US representational patterns during all conditioning phases (see Figure 2), indicating that the representation of visual information does not depend on learned associations.

##### Ventral visual structures (VVS)

Similar to V1, the rUS pattern was a significant predictor of representational patterns at all conditioning time points in ventral visual structures. Moreover, the contribution of the POI ‘Experimental Task’ was consistent across US and CS representational patterns (see Figure 2). Taken together these results indicate no effect of affective pairings on representational patterns in these visual regions.

##### Integrative regions of interest

###### Amygdala

CS patterns predicted Amygdala rUS patterns in late conditioning only, suggesting overall a slightly delayed response to learning (see Table 1). The analysis of component reactivation, however, revealed that only ‘Touch Valence’ showed an effect consistent with US data and only early in conditioning (see Figure 2).Thus, whereas the amygdala was the only ROI to recapitulate the valence of the US with appetitive and aversive at opposite poles, the effects were transient and inconsistent with the pattern of global recapitulation.

###### Ventromedial prefrontal cortex (vmPFC)

CS representational patterns in vmPFC showed clear effects of conditioning: They were not significantly predicted by vmPFC rUS pattern during early conditioning, but were during both mid and late conditioning. Examination of individual POIs indicated distinct representations of appetitive and aversive stimuli, which emerged at different time points: In mid conditioning ‘Appetitive Brush’ was reactivated, whereas in late conditioning only ‘Aversive Pressure’ showed reactivation. The results suggest that, in the vmPFC, aversive and appetitive information is carried forward in conditioning in a nonlinear fashion and with different temporal patterns.

###### Anterior cingulate cortex (ACC)

As for the vmPFC, ACC rUS could not predict ACC-CS representational patterns in early conditioning, but was highly predictive for both mid and late conditioning (see Table 1). These results indicate that ACC may represent affective associations acquired though classical conditioning. US-consistent component reactivation for ‘Experimental Task’ (similarity between all conditions within a task) by CS was observed mid and late conditioning. In addition, while ‘Aversive Pressure’ reactivation by CS data did not reach the same levels as the US, it becomes apparent that an increase in reactivation from early to mid and late conditioning is induced by conditioning (see Figure 2).

###### Insula (anterior and posterior)

Here again, no significant relationship between the anterior insula CS representational patterns and anterior insula rUS patterns was found during early conditioning, but was observed during mid and late conditioning. Similarly, the posterior insula rUS pattern was not predictive of the CS representational pattern during early conditioning, but was during mid and late conditioning (see Table 1). This suggests that global activation patterns in anterior and posterior insula represent conditioned affective associations.

The anterior insula showed clear conditioning effects, with US-consistent contribution of ‘Experimental Task’ and ‘Aversive Pressure’ in mid and late conditioning. The pattern of results suggests a bias for the anterior insula to carry forward negative hedonic information during conditioning. We also observed conditioning independent-reactivation of ‘Negative Events’ (both aversive pressure and absence of pleasurable touch) in early conditioning. On the other hand, the posterior insula only showed a US-consistent contribution of ‘Experimental Task’ in mid and late conditioning. This might represent the differences between the specific tactile and facial stimuli between the appetitive and aversive conditioning task (see Figure 2). Overall, the current data indicates some distinct patterns of reactivation in anterior and posterior insula supporting the notion that these subregions play functionally distinct roles in representing information about conditioned cues.

## Discussion

In this study we examined whether, with emotional learning, the initially neutral conditioned stimulus (CS) reactivates patterns of activation elicited by the unconditioned stimulus. We further probed whether neural population codes for the CS represent non-hedonic sensory and/or hedonic aspects of the appetitive or aversive unconditioned stimulus. In other words, we asked if what is carried forward in conditioning is the entire sensory and hedonic construct that a US encompasses, or whether only hedonic aspects of the appetitive or aversive stimulus become associated with the cue. We used representational similarity analysis (RSA) with a theoretically based version of pattern component modeling (PCM) on fMRI data to decode the content of neural representations observed over the course of aversive and appetitive classical conditioning. The data revealed that, for higher-order regions of interest, a significant amount of variance in the CS data could be explained by patterns elicited by the US, indicating US pattern reactivation by CS. Critically, this effect was only found after affective pairing had been encountered, and not prior to the learning process. By employing pattern component modeling (PCM) and determining the best combination of patterns of interest (POIs) for different ROIs, we were further able to model the distinct content of information that is represented by each brain region for the US and the degree to which it was reactivated by the CS. In primary sensory regions, the CS primarily reactivated components of the US that do not rely on learning. In contrast, several brain regions implicated in emotional learning showed reactivation of those components representing hedonic value of the US – providing evidence for a dominance of stimulus-response learning.

### Global US representation pattern reactivation by CS

We first wanted to establish whether, through conditioning, the representation of an initially neutral stimulus changes to resemble that of an inherently positive or negative one. The results from the current study suggest that, indeed, the CS reactivates patterns of US representation in all regions of interest included in the analysis. Closer examination of individual ROIs showed a more nuanced pattern of results. Visual cortex and ventral visual structures showed significant reactivation of US representational patterns across *all* experimental time points (early, mid and late conditioning). This pattern indicates that reactivation is not a result of conditioned associations but rather of experimental phase overlapping features inherent to the task (e.g., the stable visual properties of the stimuli). In contrast, amygdala, vmPFC, ACC as well as anterior and posterior insula predicted CS representational patterns from US data only *after* conditioned associations had developed. While one previous study (Onat & Büchel, 2015) has found preliminary evidence for such an effect for fear conditioning in the insula, the present study shows that this finding can be generalized to both appetitive and aversive associative learning, and to other brain regions. Thus, in this study we provide strong evidence suggesting that the basic mechanisms observed behaviorally in conditioning – that is, that the CS elicits the same response as the US (Maren, 2001; Martin-Soelch et al., 2007; Pavlov, 1927) – are also represented in the brain.

### US component reactivation by CS

After having established that, globally, the CS reactivates US representational patterns after associative learning, we next focused on the *content* of those representations. Rather than interpreting stimulus representation as a homogenous construct, we deconstructed the representation into theoretically driven patterns of interest (POIs) that allowed us to examine the aspects of the US that become attached to the CS as conditioned associations emerge. Most regions of interest included in the current analysis showed reactivation of the US representational pattern by CS. Primary visual cortex and ventral visual structures showed an experimental task-based component (POI: ‘Experimental Task’ indicating pain task vs. brush task) reactivation that did not depend on conditioning, consistent with the fact that, the visual input associated with each task did not change over the course of learning.

In contrast, component reactivation in the vmPFC and, to some degree, in the amygdala and the anterior insula, provide answers to the question of what aspects of the US become attached to the CS. A substantial body of literature has delineated complementary roles for the amygdala and vmPFC in establishing conditioning (Schoenbaum & Roesch, 2005). In the current study, in the vmPFC, representations of both appetitive and aversive touch (POIs: ‘Aversive Pressure’, ‘Appetitive Brush’) were reactivated. Of note, reactivation of appetitive touch was observed mid conditioning, while aversive touch was observed during late conditioning only.

This might indicate that the association with appetitive touch developed more quickly than with aversive pressure but that at the same time habituation to the appetitive brush is faster than to aversive pressure (Triscoli et al., 2014). First and foremost, however, the pattern of the results suggests that hedonic properties of the US are represented in the vmPFC and are reactivated during CS presentation. This finding is in line with previous studies showing that stimulus value (but not sensory properties) are represented in the vmPFC/OFC in conditioning (Chikazoe et al., 2019; Lim et al., 2013; McNamee et al., 2013). The amygdala was the only region where the representation of the hedonic experience of touch valence (POI. ‘Touch Valence’) was reactivated, rather than non-specific sensory input or aversive pressure. Yet the contribution of ‘Touch Valence’ was consistent with the US data early in conditioning only. This pattern of results shows that the amygdala represents appetitive and aversive associations as polar ends on a valence spectrum. This finding is consistent with recent evidence in mice and humans for spatially segregated antagonistic valenced neurons and, in humans, coding of a spectrum of valence in the amygdala (Jin et al., 2015; Kim et al., 2016). It should be highlighted, however, that in the current task this effect did not seem to emerge with conditioning but was rather inherent to the processing of the facial stimulus alone.

Interestingly, while the US representations in anterior and posterior insula showed large overlap, a distinctive role for each of the two regions emerged for the CS during conditioning. While both anterior and posterior insula activation showed a contribution of the task-based model (POI: ‘Experimental Task’) that contrasts appetitive and aversive conditioning tasks in general, the anterior insula further showed reactivation of components representing negatively valenced information. Much research has been done on the role of the insula and its subregions with somewhat inconsistent results. A recent meta-analysis (Kurth et al., 2010) revealed that most functions investigated have been associated with activation in the anterior portion, especially emotional processing. Only sensorimotor processing has been exclusively mapped onto the posterior insula while pain processing has been shown to involve the entire insula (Craig, 2002). In the light of these meta-analysis findings, the current results of reactivation of negative information representation in the anterior insula, especially without reactivation of non-specific sensory input (reactivation POI: ‘Non-specific Touch’) strongly suggest reactivation of hedonic but not sensory US components. In contrast, the task-based (reactivation POI: ‘Experimental Task’) in the posterior insula could be explained by the differences in sensory input between appetitive and aversive tasks, i.e. female faces paired with appetitive brush and male faces paired with aversive pressure respectively. In summary, after demonstrating US component reactivation by CS with conditioning, the current findings show that several brain regions that have been previously associated with conditioning are biased to reactivate components that carry hedonic information instead of purely tactile information. Thus, the pattern of results observed here suggests that what is carried forward in conditioning and becomes associated with the CS is primarily the affective attachment with the stimulus rather than the pure sensory experience. In other words, when we are exposed to a conditioned stimulus we re-experience the pleasant or unpleasant feelings elicited by the US rather than the sensory stimulation. Taken together, the current analysis provides support for the notion of stimulus-response conditioning in which a CS reproduces the unconditioned response rather than the perceptual experience of the US.

In conclusion, we demonstrated for the first time that, in conditioning, a conditioned stimulus reactivates the pattern of activation initially elicited by the unconditioned stimulus in several brain regions. The results further show that it is primarily the hedonic components of the initial experience, rather than discriminative sensation, that is reproduced when we encounter a cue that predicts it. Thus, when we encounter a cue signaling pain or pleasure, what we carry forward is the emotional meaning we attach to it rather than the sensory experience.

## Materials and Methods

### Participants

Data from 71 young, healthy participants (age: 21.1 ± 2.8 years, 41 females) was included in the analysis. Initially, 107 participants were recruited from Cornell University to participate in a brain imaging study of appetitive and aversive classical conditioning tasks. A number of participants had to be excluded for the following reasons: 20 participants had missing data (imaging run, stimulus onset files, motion correction files) while multi-echo preprocessing described below failed for 16 participants. All participants gave written, informed consent and had normal or corrected-to-normal vision. Participants were pre-screened for a history of anxiety and depression as well as other psychopathology, epilepsy and brain surgery in addition to general suitability for fMRI data collection. Pre-screening was followed up in person by an additional interview to ensure inclusion criteria were met. Due to the fact that this study was conducted as part of a larger research program, all participants were genotyped.

The neural representation of the appetitive and aversive US as well as the development of the PCM derivative and other methodical details have been described in (Kryklywy, Ehlers, et al., 2021).

### Materials

#### Stimulus and apparatus

Six faces were chosen from the Karolinska directed emotional faces, comprising three male and three female exemplars each with a neutral expression (Goeleven, De Raedt, Leyman, & Verschuere, 2008). These faces were used as the conditioned stimuli (CS) in a classical conditioning paradigm. The (US) consisted of either an aversive pressure delivered to the right thumb, or an appetitive brush stroke to the participant’s forearm. Aversive pressure stimuli were delivered using a custom designed hydraulic device, similar to those used in previous studies (Giesecke et al., 2004; López-Solà et al., 2010), capable of transmitting controlled pressure to 1 cm^2^ surface placed on the subjects’ right thumbnail. In individual calibration sessions, it was ensured that the pressure intensity was aversive but not excessively painful. Appetitive brush strokes were manually applied to the left forearm lasting ∼4s. Individual subjective responding to brush stimuli were recorded in a separate session prior to all experimental scanning, with only participant who responded positively to the manipulation invited to participant in the scanning session.

### Procedure

#### Stimulus ratings

As a measure of subjective stimulus assessment and conditioning, participants were asked to rate the likeability and trustworthiness of the faces used as CS+ and CS- stimuli on a scale from 1-100 (1) before and (2) after conditioning as a measure of conditioning. CS Discrimination scores ([CS+ minus CS-]) before and after conditioning were calculated and compared using t-tests. Due to technical difficulties, stimulus ratings were only available for 69 of the 71 participants included in the analysis.

#### Experimental tasks

While undergoing functional MR scanning, participants completed two unique conditioning tasks with nearly identical structure modeled after Visser and colleagues (2015). These tasks differed from each other only in the nature of the tactile unconditioned stimulus (US; see above), and the gender of the face stimuli. In each task, participants completed seven CS- only blocks interleaved with six CS- US paired blocks (see Figure 2a). Single blocks of either the CS- only or the CS- US pairing entailed one presentation of each of the three male or female face stimuli used in that task. The order of the CS+ and the CS- faces was randomized within each CS- US block. Individual trials started with an initial fixation period (19500 ms) followed by the presentation of a face stimulus (4000 ms). The fixed and long interstimulus interval was included in the experimental design to reduce intrinsic noise correlations (Visser et al., 2013). During CS- only trials, all faces were presented without tactile stimulation (see Figure 1d). During CS- US paired trials, two of three facial stimuli were paired with tactile stimulation, thus creating two CS+ and one CS- face stimuli (see Figure 2b). The US was delivered from the midpoint after the visual stimulus presentation (2000 ms post-onset), and remained for the duration of the visual presentation (2000 ms). Face pairings were randomly assigned for each participant but held constant across the duration of the experiment.

### MRI acquisition and preprocessing

#### Acquisition

Scanning was conducted on a 3 Tesla GE Discovery magnetic resonance scanner using a 32-channel head coil at Cornell University. For each subject, a T1-weighted MPRAGE sequence was used to obtain high-resolution anatomical images (repetition time (TR) = 7 ms, echo time (TE) = 3.42 ms, field of view (FOV) 256 x 256 mm slice thickness 1 mm, 176 slices). The functional tasks were acquired with the following multi-echo (ME) EPI sequence: TR = 2000 ms, TE1 = 11.7 ms, TE2 = 24.2 ms and TE3 = 37.1 ms, flip angle 77°; FOV 240 x 240 mm. A total of 102 slices was acquired with a voxel size of 3 x 3 x 3 mm. Pulse and respiration data were acquired with scanner-integrated devices.

#### Preprocessing

Multi-echo independent component analysis (ME-ICA, meica.py version 3.2 beta1) was used to denoise the multi-echo fMRI data. An optimally combined (OC) dataset was generated from the functional multi-echo data by taking a weighted summation of the three echoes, using an exponential T2* weighting approach (Posse et al., 1999). Multi-echo principal components analysis was first applied to the OC dataset to reduce the data dimensionality. Spatial independent component analysis (ICA) was then applied and the independent component time-series were fit to the pre-processed time-series from each of the three echoes to generate ICA weights for each echo. These weights were subsequently fitted to the linear TE-dependence and TE-independence models to generate F-statistics and component-level κ and ρ values, which respectively indicate blood-oxygen-level-dependent (BOLD) and non-BOLD weightings (Kundu et al., 2012). The κ and ρ metrics were then used to identify non-BOLD-like components to be regressed out of the OC dataset as noise regressors (Kundu et al., 2013).

### Functional imaging analyses

#### Regions of interest

To assess tactile (aversive pressure, appetitive brush) and hedonic representations in neural patterns, eight bilateral regions of interest (ROIs) were generated from the standard anatomical atlas (MNI_caez_ml_18) implemented in the Analysis of Functional NeuroImages (AFNI) software package (Cox, 1996): primary somatosensory cortex (S1), primary/secondary visual cortex (V1) were selected as the primary sites of tactile and visual information respectively. In addition, ventral visual structures (VVS) were chosen due to their role in visual classification (Kanwisher et al., 1997; Kravitz et al., 2013). Amygdala, ventromedial prefrontal cortex (vmPFC), anterior cingulate cortex (ACC) and insula were further selected for their hypothesized roles in affect processing (Anderson & Phelps, 2002; Craig, 2002; Winecoff et al., 2013) and pain representations (Kragel et al., 2018; Orenius et al., 2017). The insula was further divided into an anterior and posterior portion due to its functional and anatomical subdivisions (Nieuwenhuys, 2012) for a total of eight ROIs.

#### Representational similarity analysis

Data analysis of the fMRI data was conducted using Analysis of Functional NeuroImages (AFNI) software (Cox, 1996). Regressor files of interest were generated for all individual trials across the experiment, modelling the time course of each stimulus presentation during each run (36 total events). The relevant hemodynamic response function was fit to each regressor to perform linear regression modeling. This resulted in a β coefficient and t value for each voxel and regressor. To facilitate group analysis, each individual’s data were transformed into the standard brain space of Montreal Neurological Institute (MNI).

In order to identify the representational pattern elicited by the experimental stimuli, representational similarity analysis (RSA) was performed using the Python package PyMVPA (Hanke et al., 2009). For each participant, in each ROI the spatial pattern of β weights in response to each experimental condition or event was correlated with the pattern of activation in response to all other events. This step was performed separately for each ROI. Thus, pairwise Pearson coefficients for all experimental events of a single ROI resulted in a similarity matrix containing correlations for all CS- US combinations for all trials for each participant. Fischer transformations were performed on all similarity matrices to allow comparisons between participants.

#### Pattern Component Modeling

In order to characterize the content of CS (and US) representation in key regions of interest, we developed a theory-guided implementation of Pattern Component Modeling (PCM) (Diedrichsen et al., 2018; Kriegeskorte & Kievit, 2013; Kryklywy, Ehlers, et al., 2021; Kryklywy, Forys, et al., 2021). The details are described in Kryklywy, Ehlers, et al., 2021. In brief, we created 13 patterns of interest (POIs) to represent dissociable correlation patterns that would be observed in the experimental data if it would contain perfect representation of distinct theoretically-derived constructs (Fig 1C). POIs were constructed for 1) Experimental Task, 2) Non-Specific Touch, 3) Specific Touch, 4) Appetitive Brush, 5) Aversive Pressure, 6) Touch Valence, 7) Positive Events, 8) Negative Events, 9) All Valence, 10) Salience, 11) Face Stimulus, 12) Violation of Expectation and 13) Temporal Adjacency (see Figure 1c). In order to determine the POI combinations that best explained the observed correlation in the US data in each ROI, a Bayesian Information Criterion (BIC) analysis and multiple regression implemented in our R package ‘PCMforR’ (Kryklywy, Forys, et al., 2021) were conducted. A reconstructed US (rUS) pattern was built from identified POIs in order to reduce noise. The rUS pattern was first used as predictor for the CS pattern in early, mid and later conditioning to determine the global reactivation of US patterns by CS- only data. Subsequently, the US identified POIs were used as predictors in Bayesian linear models for both rUS and CS data in order to compare the contribution (beta weight) of each POI between rUS and CS data. For that purpose the R package ‘BayesFactor’ (Morey et al., 2018) was used. The Bayesian linear model was estimated with 1,000,000 iterations allowing us to extract mean beta weights for each POI and their 95 % credible intervals (CrIs). In order to determine whether the contribution of each POI to the CS data is comparable to that of the US data, we adapted an approach developed to assess the robustness of replications (LeBel et al., 2018) that has recently also been employed in a Bayesian framework (Kuhn et al., 2021). While we are not comparing replication attempts, we have adapted the measure of consistency described previously (LeBel et al., 2018) in such a way that consistency between US and CS data is assumed when the beta weight point estimate obtained from CS data of any given POI is included in the credible interval of the beta weight obtained from US data for the same POI.

## Acknowledgements

This research was supported by Canadian Institutes for Health Research (CIHR) Operating Grant #491746 and a grant from the Leaders Opportunity Fund from the Canadian Foundation for Innovation (FAS#: F13-03917) as well as a CIHR New Investigator Salary Award, and a Michael Smith Foundation for Health Research Scholar award to R.M. Todd. J.H. Kryklywy is supported by a Natural Sciences and Engineering Research Council of Canada (NSERC) fellowship award (PDF-532611-2019).

## Author Contributions

M.R. Ehlers: conceptualization, methodology, formal analysis, data curation, writing – original draft, visualization, project administration

J.H. Krkylywy: conceptualization, methodology, software, formal analysis, data curation, writing – original draft, visualization

A.O. Beukers: methodology, software, formal analysis, writing – review & editing

S.R. Moore: conceptualization, software, project administration

B.J. Forys: formal analysis, data curation, writing – review & editing

A.K. Anderson: conceptualization, methodology, resources, writing – review & editing, supervision, funding acquisition

R.M. Todd: conceptualization, methodology, resources, writing – original draft, supervision, funding acquisition

